# Efficient and highly amplified imaging of nucleic acid targets in cellular and histopathological samples with pSABER

**DOI:** 10.1101/2023.01.30.526264

**Authors:** Sahar Attar, Valentino E. Browning, Yuzhen Liu, Eva K. Nichols, Ashley F. Tsue, David M. Shechner, Jay Shendure, Joshua A. Lieberman, Shreeram Akilesh, Brian J. Beliveau

## Abstract

*In situ* hybridization (ISH) is a powerful tool for investigating the spatial arrangement of nucleic acid targets in fixed samples. ISH is typically visualized using fluorophores to allow high sensitivity and multiplexing or with colorimetric labels to facilitate co-visualization with histopathological stains. Both approaches benefit from signal amplification, which makes target detection effective, rapid, and compatible with a broad range of optical systems. Here, we introduce a unified technical platform, termed ‘pSABER’, for the amplification of ISH signals in cell and tissue systems. pSABER decorates the *in situ* target with concatemeric binding sites for a horseradish peroxidase-conjugated oligonucleotide which can then catalyze the massive localized deposition of fluorescent or colorimetric substrates. We demonstrate that pSABER effectively labels DNA and RNA targets, works robustly in cultured cells and challenging formalin fixed paraffin embedded (FFPE) specimens. Furthermore, pSABER can achieve 25-fold signal amplification over conventional signal amplification by exchange reaction (SABER) and can be serially multiplexed using solution exchange. Therefore, by linking nucleic acid detection to robust signal amplification capable of diverse readouts, pSABER will have broad utility in research and clinical settings.

## Introduction

Since its introduction by Pardue and Gall^1^ in the late 1960s, *in situ* hybridization (ISH) has been widely use in research and clinical settings to determine the abundance and subcellular location of nucleic acid targets. ISH relies on the formation of heteroduplexes between target RNA and DNA strands present in fixed cellular and tissue samples and complementary probe strands that are added during a hybridization reaction. Once completed, the results of the hybridization reaction can be visualized using transmitted light or fluorescent microscopy depending on the type of labeling strategy used. Common applications for ISH include targeting intervals of genomic DNA to assay chromosome copy number in cytogenetic clinics^2^, the quantitative profiling of single-cell RNA expression profiles^3^, the investigation of how chromosomes are spatially organized within the nucleus^4^, and the visualization of gene expression patterns in whole mount tissue specimens^5^.

Considerable technical development has occurred to improve the sensitivity of ISH and to broaden its applications. Among these, notable developments include the transition from radioactive to fluorescent labels^6–8^, the use of recombinant^9^ and synthetic^3,10–13^ DNA as a source of probe material, and the introduction of massively multiplexed imaging approaches that rely on iterative rounds of hybridization and imaging in the same sample^14–16^. Another set of technical development has focused on strategies to amplify the signal of ISH, which has particular benefits in challenging sample types such as formalin-fixed, paraffin-embedded (FFPE) tissue sections, thick frozen tissue sections, and whole mount tissue samples^17,18^. Broadly, these signal amplification approaches fall into two categories: 1) Assembly-based technologies in which DNA and/or protein species are directed to the site(s) of ISH in order to create an *in situ* scaffold capable of recruiting many directly labeled molecules to increase the overall label density at the target site. Examples of assembly-based approaches include the assembly of branched DNA (bDNA)^19,20^ and LNA^21^ structures, hybridization chain reaction (HCR)^22^, and ClampFISH^23,24^; 2) Enzymatic approaches in which specialized probe sets recruit enzymes to catalyze the formation signal amplification structures *in situ*. Examples of enzymatic approaches include the catalyzed reporter deposition (CARD) technique that uses the horseradish peroxidase (HRP) enzyme to locally deposit detectable biotin molecules^25^, methods that recruit HRP or alkaline phosphatase for colorimetric detection via haptenized probes in whole-mount embryos^26^, and rolling circle amplification (RCA)^27^.

Assembly-based and enzymatic signal amplification approaches can strengthen signals by >100x, making it easier to detect analytes in thick and noisy sample types. These methods even make it possible to image labeled specimens on low numerical aperture imaging systems with lower photon collection efficiencies. However, as these approaches require that the signal amplification structure be created *in situ*, they can be difficult to troubleshoot, particularly for researchers who are performing ISH itself for the first time. Moreover, many of these approaches rely on costly or proprietary reagents, which adds to the difficulty of establishment and implementation. The recently developed signal amplification by exchange reaction (SABER) technique introduced by Kishi and colleagues^28^ abrogates these issues, as signal amplification in SABER is achieved by the *in vitro* addition of long, concatemeric ssDNA tails to the probe molecules via the primer exchange reaction (PER)^29^ prior to their addition to the fixed sample. As the creation of the signal amplification structure occurs *in vitro*, researchers can evaluate the results of the PER reaction and troubleshoot as needed prior to executing time consuming ISH experiments with precious samples. In addition to this useful feature, SABER has many strong attributes as a signal amplification approach, including low-cost ($1–5 per sample) simple and robust multiplexing via solution exchange^28^.

While SABER has features that may accelerate its adoption, it still has limitations. For instance, SABER can achieve ∼450x signal amplification when it employs iterative rounds of branched assembly *in situ*^28^, but to date that approach has only be shown for imaging one target at a time and has only been used on cultured cells, and the unbranched form of SABER that supports multicolor imaging in cells and tissues only provides a more modest ∼5–15x amplification^28^. SABER also has yet to be combined with colorimetric detection methods, which facilitate co-visualization with histopathological stains such as hematoxylin and eosin. Here, we introduce ‘peroxidase-SABER’ (pSABER), a robust and cost-effective strategy for boosting the signal of SABER by a further 20-fold and expanding its readout modalities to include colorimetric detection. We demonstrate the effectiveness of colorimetric and fluorescent pSABER in fixed cells and FFPE tissue sections and showcase the ability of pSABER to perform highly amplified and multiplexed imaging via solution exchange. We anticipate that its simplicity and its seamless integration into the existing SABER toolkit will lead to the adoption of pSABER as a workhorse technique for *in situ* nucleic acid detection in research and clinical settings.

## Results

### Programing enzymatic signal amplification with pSABER

The pSABER approach builds on the established, unbranched SABER workflow. First, probe sets or “readout oligos” complementary to a common exogenous binding site encoded in complex oligo probe libraries are extended *in vitro* by the primer exchange reaction (PER)^28,29^, resulting in the addition of long concatemers of a reiterated 10 nt PER sequence (Fig. 1). These concatemers are then applied to the sample and primary *in situ* hybridization is performed (and secondary hybridization with extended readout oligos in the case that readout oligos are being used). To detect the hybridized concatemers, the sample is then hybridized with a 20 nt oligo covalently conjugated to HRP that is complementary to a dimer of the reiterated PER sequence (Fig. 1). Finally, the sample is incubated with H_2_O_2_ and an HRP substrate to facilitate the localized deposition of a variety of colorimetric or fluorescent labels that can be detected by microscopy (Fig. 1).

**Fig. 1.**
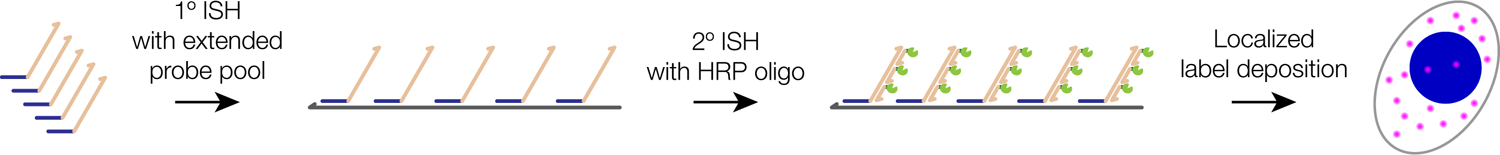
Schematic overview of pSABER. *In vitro* extended primary oligo probes are applied to samples and hybridized *in situ* to DNA or RNA targets, facilitating the recruitment of a secondary oligo conjugated with horseradish peroxidase (HRP) that can locally catalyze the deposition of colorimetric or fluorescent labels.

### pSABER enables amplified colorimetric detection

In order to validate the pSABER approach and demonstrate its compatibility with colorimetric detection, we designed a series of RNA and DNA ISH experiments against well-established molecular targets. We first performed an RNA ISH experiment using a probe set consisting of 83 oligos targeting the *ADAMTS1* mRNA in human K29Mes cells that had been shifted to 37°C prior to fixation to induce *ADAMTS1* expression^30^. After primary hybridization, the sample was developed with 3,3’-Diaminobenzidine (DAB) in the presence of H_2_O_2_ prior to hematoxylin staining and mounting. Upon visualization by routine brightfield microscopy, both robust cytoplasmic DAB staining consistent with single-molecule RNA FISH^3^ and strong nuclear puncta corresponding to sites of induced transcription^30^ were observed, neither of which was visible in the no primary probe negative control (Fig. 2a). To test the applicability of pSABER to more challenging DNA ISH applications, we performed experiments targeting the human alpha satellite repeat DNA region in K29Mes cells (Fig. 2b) as well as a 200 kb single-copy chromosomal region on Xp22.32 (hg38 chrX:5400000–5600000) using a previously validated complex oligo library and an accompanying readout probe^31^ on primary human metaphase chromosome spreads (Fig. 2c). In both cases, brightfield microscopy revealed strong, punctate staining in the expected patterns specific to DNA, whereas no appreciable staining was observed in the corresponding no primary probe controls (Fig. 2b,c). Finally, we examined the ability of colorimetric pSABER to detect targets in FFPE tissue sections, which represents the main specimen format in clinical and research pathology laboratories worldwide. We first performed RNA ISH in deidentified human kidney FFPE sections using a probe set consisting of 6 oligos targeting the internal transcribed spacer 1 (ITS1) region of the 47S pre-rRNA that is localized exclusively in the nucleolus^32^. After colorimetric development, we observed tightly localized signal inside of nearly every nucleus in the section, with the expected 1–2 signals per nucleus being observed (Fig. 2d). We also verified colorimetric RNA pSABER can detect RNA originating from a non-repetitive locus in FFPE specimens by performing RNA ISH with a probe set consisting of 83 oligos targeting the *UMOD* mRNA in deidentified human kidney sections. Using pSABER we detected a very specific pattern of UMOD transcript localization in cells of the thick ascending loop of Henle (TALH) (arrowheads, Fig. 2e, center panel), but not in cells of the adjacent macula densa (MD), a specialized structure in the vicinity of the kidney glomerulus-tubule junction. This localization pattern exactly matched UMOD protein expression as determined by immunohistochemistry in the human protein atlas^33^. No detectable staining was observed in the corresponding no primary probe controls for the kidney FFPE experiments (Fig. 2d,e). These experiments further validated pSABER for *in situ* localization of mRNA molecules in clinically relevant samples.

**Fig. 2.**
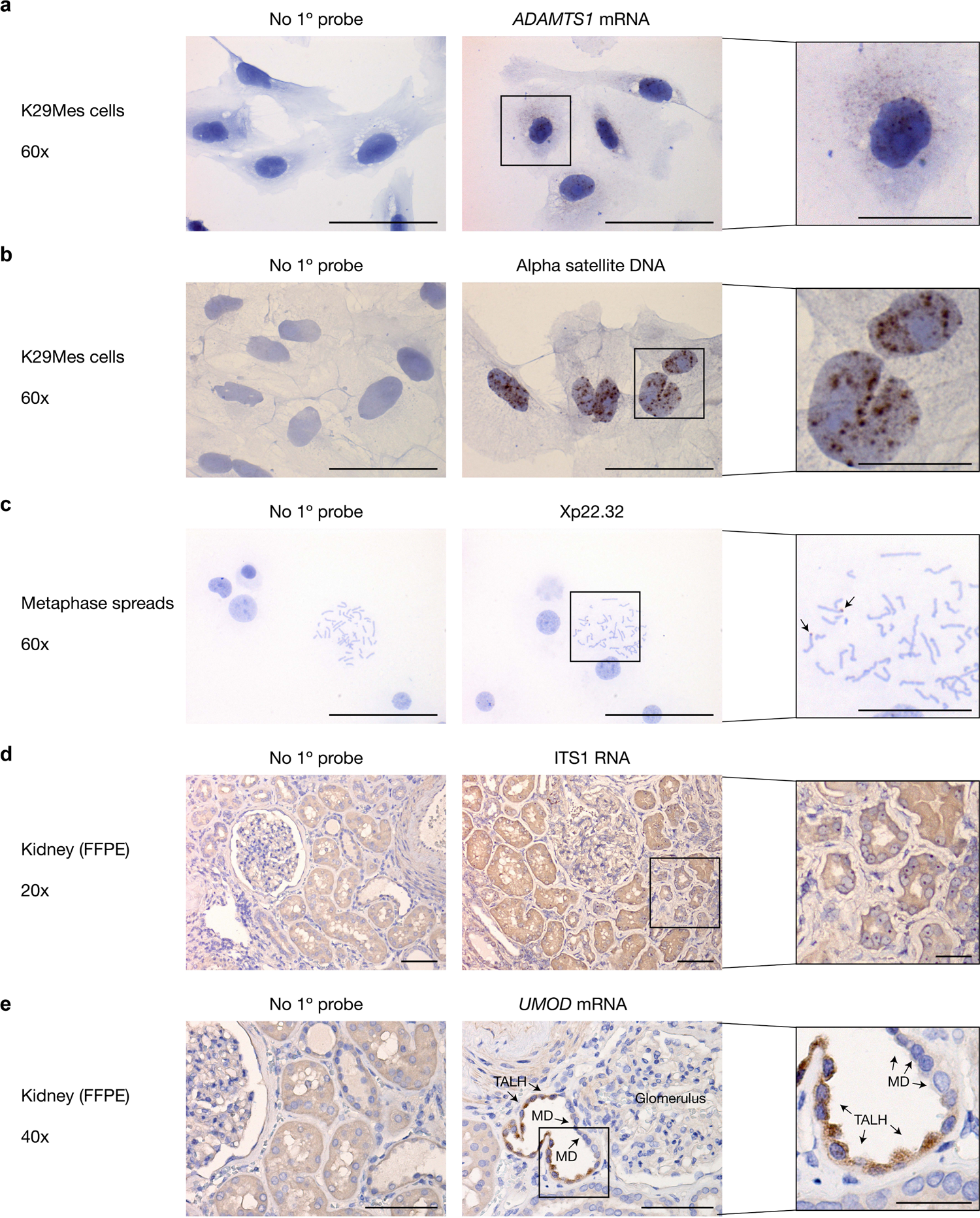
pSABER enables colorimetric detection of nucleic acid targets. **a**, Visualization of the *ADAMTS1* mRNA in human K29Mes cells. **b**, Visualization of alpha satellite DNA in mRNA in human K29Mes cells. **c**, Visualization of a 200 kb genomic interval within Xp22.32 on primary human metaphase chromosome spreads. **d**, Visualization of ribosomal RNA internal transcribed spacer 1 (ITS1) region in deidentified primary human paraffin fixed, formalin embedded (FFPE) kidney tissue sections. **e**, Visualization of the *UMOD* mRNA in deidentified primary human FFPE kidney tissue sections. A no primary probe negative control image processed in parallel to each sample is shown to the left of each pSABER image. Scale bars, 20 µm (insets) or 50 µm (fields of view). MD = macula densa. TALH = thock ascending loop of Henle.

### Enhanced signal amplification with fluorescent pSABER

HRP can also catalyze covalent deposition of fluorophore-conjugated tyramides in a variety of specimen types. Therefore, we next set out to determine whether pSABER could generate amplified fluorescent signals *in situ* via the HRP-catalyzed deposition of fluorophore-conjugated tyramides at the sites of primary probe binding. To this end, we performed a series of fluorescent RNA pSABER experiments in EY.T4^34^ mouse embryonic fibroblast cells using fluorescein-tyramide (Methods). During the development step, we independently targeted each of three RNA species with distinct expected spatial signal patterns: 1) the 47S rRNA ITS1 region that is localized exclusively in the nucleolus^32^ (Fig. 3a); 2) the 7SK small nuclear RNA which is broadly localized within the nucleoplasm and enriched in nuclear speckles, but excluded from nucleoli^35^ (Fig. 3b); the *Cbx5* mRNA which localizes predominantly in the cytoplasm^28^ (Fig. 3c). In all three cases, using confocal microscopy, we observed the expected staining patterns, whereas we saw no detectable signal in the no primary probe negative control (Fig. 3b). We also validated that fluorescent pSABER can faithfully visualize nucleic acid targets in FFPE specimens by performing an experiment targeting ITS1 in deidentified human kidney FFPE tissue sections. This revealed the expected pattern of strong nucleolar staining (Fig. 3e) that matched the signal seen with colorimetric DAB development (Fig. 2d) and that was not observed in the matched no primary probe negative control (Fig. 3f).

**Fig. 3.**
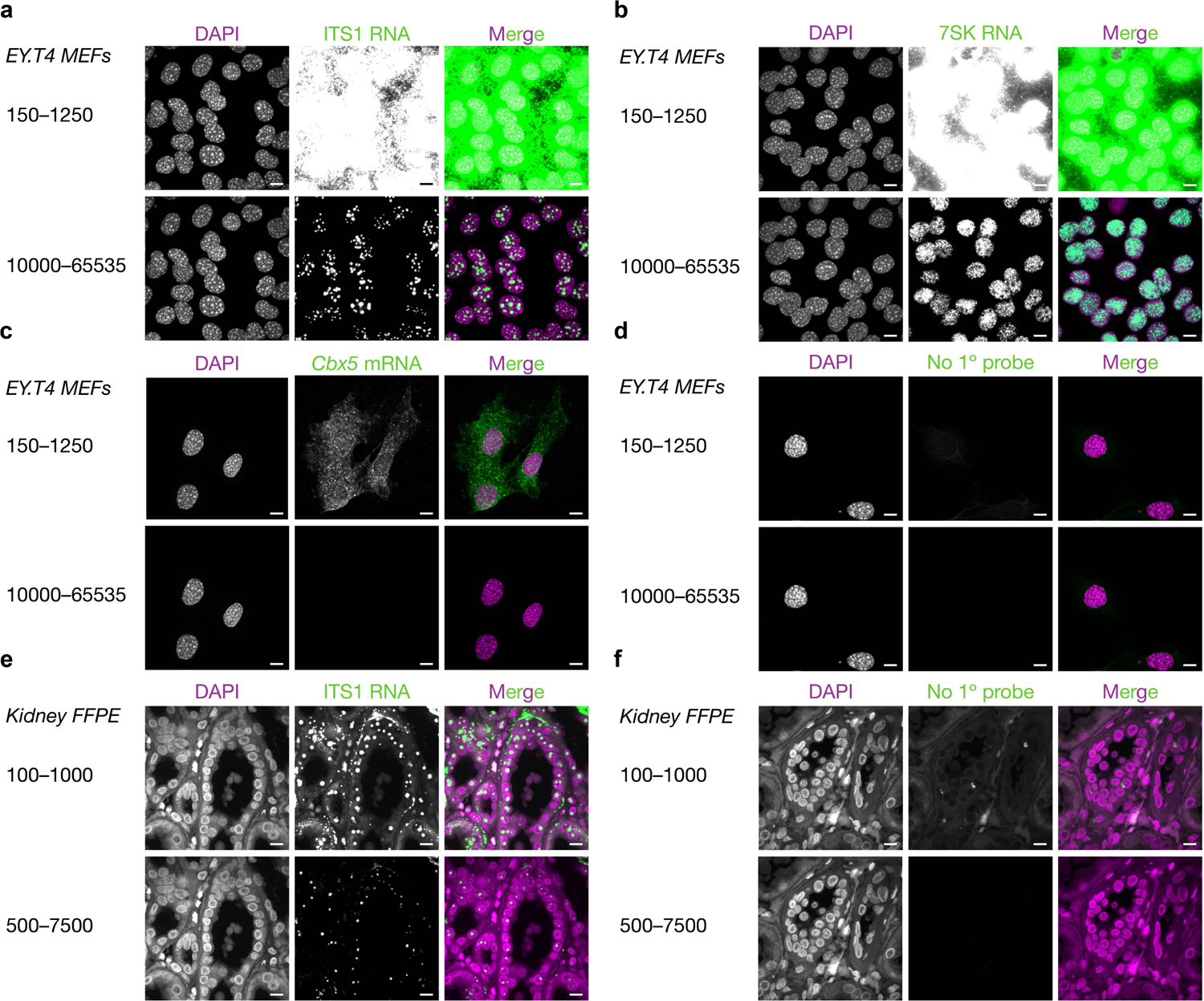
Fluorescent pSABER visualizes nucleic acid targets in cells and tissues. **a**, DNA (DAPI, left), fluorescent pSABER targeting the ribosomal RNA internal transcribed spacer 1 (ITS1) region performed in EY.T4 mouse embryonic fibroblasts (MEFs) (center), and a composite image (right). **b**, DNA (DAPI, left), fluorescent pSABER targeting the 7SK RNA performed in EY.T4 MEFs (center), and a composite image (right). **c**, DNA (DAPI, left), fluorescent pSABER targeting the *Cbx5* mRNA performed in EY.T4 MEFs (center), and a composite image (right). **d**, DNA (DAPI, left), a no primary probe control for pSABER performed in EY.T4 MEFs (center), and a composite image (right). **e**, DNA (DAPI, left), fluorescent pSABER targeting ITS1 performed in deidentified primary human paraffin fixed, formalin embedded (FFPE) kidney tissue sections (center), and a composite image (right). **f**, DNA (DAPI, left), a no primary probe control for pSABER performed in deidentified primary human FFPE kidney tissue sections (center), and a composite image (right). Each set of images is presented at the indicated high (top) and low (bottom) contrast settings to allow for visual comparisons between signals occupying different portions of the 16-bit dynamic range (0–65535). Images are maximum projections in Z. Scale bars, 10 µm.

After verifying that pSABER is readily compatible with fluorescent detection, we next set out to quantify the degree of signal amplification provided by pSABER. In particular, we aimed to measure how much, if any, additional amplification pSABER provides over the conventional version of the SABER technique. To this end, we performed a paired series of ISH experiments targeting the centromeric mouse minor satellite repeat in EY.T4 MEF cells in which the same pool of PER-extended probe was hybridized and visualized either via the recruitment of an ATTO 488 labeled oligo (SABER) or via the recruitment of an HRP-oligo followed by development with fluorescein-tyramide (pSABER) (Fig. 4a). Both samples were acquired using matched illumination and camera exposure settings. These settings were optimized to utilize the full dynamic range of the camera without significant amounts of pixel saturation (Methods), allowing a direct and quantitative comparison of the signal intensities of each sample (Fig. 4b). Following automated segmentation nuclei and minor satellite FISH puncta (Methods), pSABER was observed to produce a distribution of peak signal intensity per puncta values whose median was 25.1x greater than the corresponding SABER distribution (Fig. 4c). A similar fold-enhancement (21.3x) was observed when comparing the means of the distributions (Fig. 4c). From these data, we conclude that fluorescent pSABER provides substantial signal amplification (>20x) relative to conventional SABER. Furthermore, as SABER was reported to produce ∼10x signal amplification relative to unamplified FISH at similar targets^28^, we estimate that fluorescent pSABER provides >200x amplification relative to unamplified FISH.

**Fig. 4.**
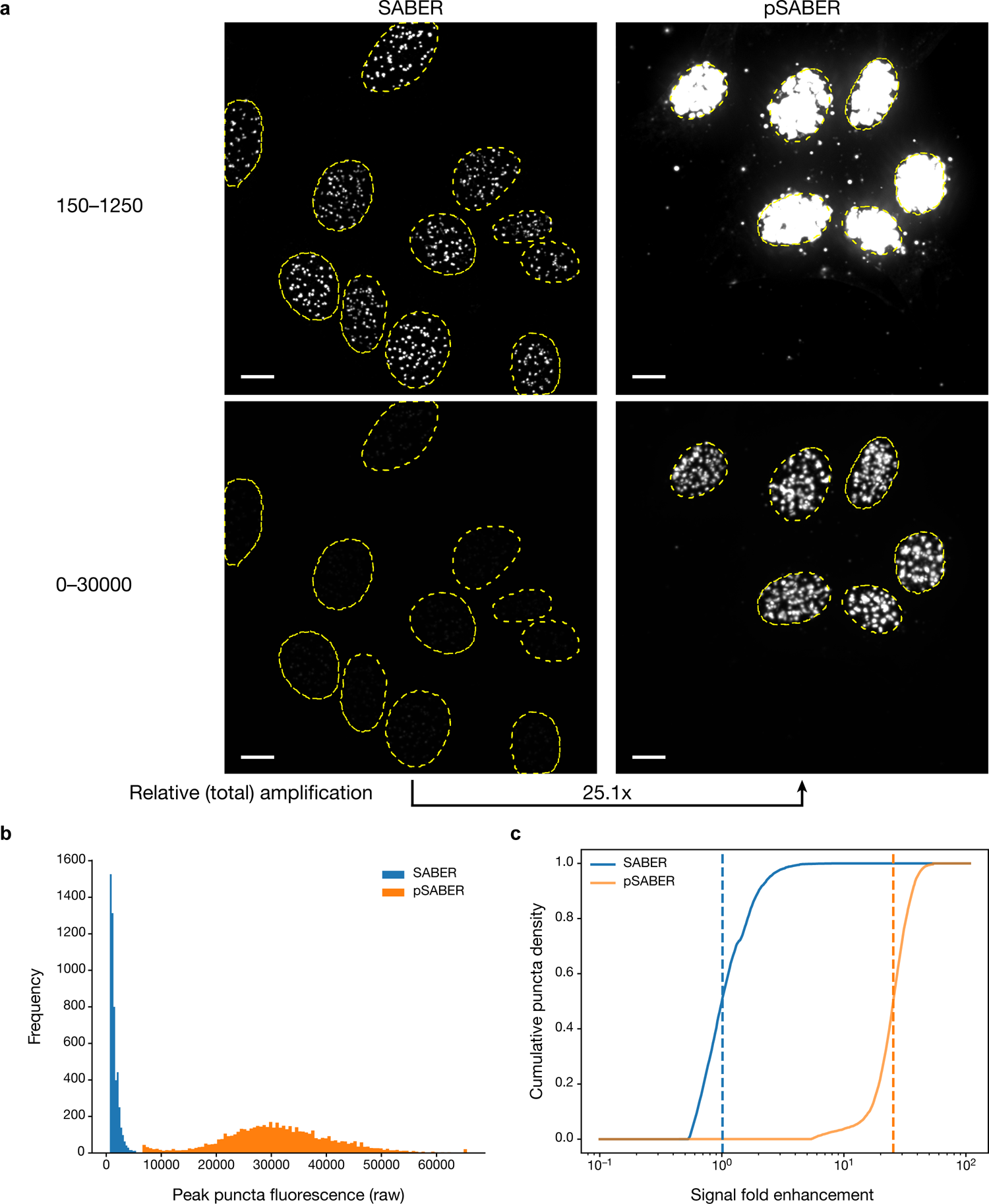
pSABER robustly amplifies fluorescent signals *in situ*. **a**, Representative images of DNA FISH targeting the mouse minor satellite repeat in EY.T4 mouse embryonic fibroblasts visualized by SABER (left) or pSABER (right). Outlines of nuclei segmented using the DAPI signal (yellow, dashed line) are overlaid on all images. Each image is presented at the indicated high (top) and low (bottom) contrast settings to allow for visual comparisons between signals occupying different portions of the 16-bit dynamic range (0–65535). **b**, Distributions of the peak pixel intensity of each segmented puncta (see Methods). **c**, Normalized empirical cumulative distributions of the peak pixel intensity of each segmented puncta showing fold-enhancement over SABER, with vertical dashed lines corresponding to medians. A similar value of fold-enhancement was obtained comparing means (25.1x enhancement, medians; 21.3x enhancement, means). *n*_SABER_ = 5,171 puncta; *n*_pSABER_ = 5,770 puncta. Scale bars, 10 µm.

### Exchange-pSABER for multiplexed detection with amplification

A powerful feature of SABER is its ability to sequentially visualize multiple targets via the programmable displacement of bound fluorescently labeled oligos and the subsequent hybridization of labeled oligos to distinct targets bearing orthogonal sequence extensions^28^. We reasoned that fluorescent pSABER could also harness this property to enable multiplexed and highly amplified imaging via solution exchange of HRP-oligos between subsequent rounds of signal development (Fig. 5a). To test this idea, we designed a 3-color imaging experiment targeting the *Cbx5* mRNA, the 7SK small nuclear RNA, and the ITS1 region of the 47S rRNA. In this experiment, all primary probes were first co-hybridized in a single overnight step. Next, we added the HRP-oligo complementary to the “p27” SABER extension borne by the *Cbx5* probe set and developed this with Cy5-tyramide (Fig. 5b, top). We then removed the bound HRP-oligo by solution exchange into formamide buffer^28^ (Methods), allowing us to perform a second round of hybridization of with the HRP-oligo, but this time complementary to the “p30” extension borne by the 7SK probe set and developed with Cy3-tyramide (Fig. 5b, middle). In the final cycle, we removed the bound HRP-oligo via solution exchange and performed a third round of hybridization and development at ITS1 with fluorescein-tyramide (Fig. 5c, bottom).

**Fig. 5.**
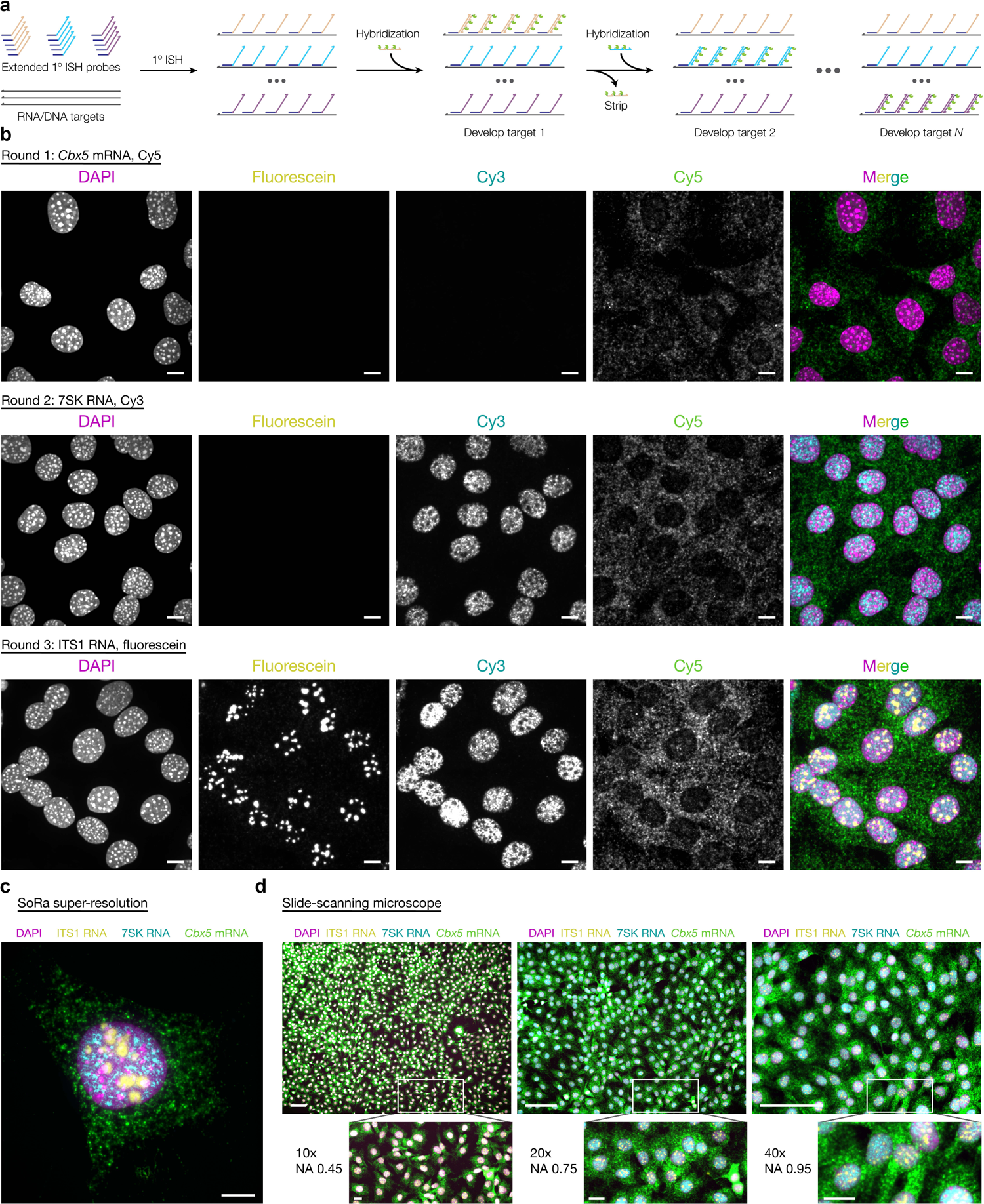
Exchange-pSABER enables highly amplified and multiplexed imaging. **a**, Schematic overview of Exchange-pSABER depicting sequential rounds of hybridization with an HRP-oligo, development, displacement, and re-hybridization. **b**, 3 color Exchange-pSABER experiment in EY.T4 mouse embryonic fibroblasts targeting the *Cbx5* mRNA (round 1, Cy5, top), the 7SK RNA (round 2, Cy3, middle), and the ribosomal RNA internal transcribed spacer 1 (ITS1) region (round 3, fluorescein, bottom). **c**, A composite image of the 3-color Exchange-pSABER experiment depicted in panel b taken using a SoRA CSU-W1 super-resolution spinning disc microscope. **d**, Composite images of the 3-color Exchange-pSABER experiment depicted in panel b taken using an automated slide-scanning microscope using 10x (left), 20x (center), and 40x (right) air objectives. Images are maximum projections in Z (a–c) or single Z slices (d). Scale bars, 10 µm (a–c, fields of view), 20 µm (d, insets), or 100 µm (d, fields of view).

Visualization after each round of development revealed the sequential accumulation of amplified signal patterns at the three targets, with the fluorescent tyramides added during previous rounds remaining stable and prominently visible even during later rounds (Fig. 3b). After the three rounds, we observed that signal associated with each RNA target was properly localized to the correct subcellular location with minimal spatial overlap even when visualized using “SoRa” super-resolution spinning disc confocal microscopy (Fig. 3c). Moreover, we found that we could observe the amplified, multiplexed staining pattern even when using lower numerical aperture air objectives on an automated slide scanning microscope at 40x, 20x and even 10x magnifications (Fig. 4d), whereas we were essentially unable to detect conventional SABER using the same system (Supplementary Fig. 1).

## Discussion

The pSABER approach sits conveniently upon the established PER/SABER platform to expand its capabilities to colorimetric detection and highly amplified (>200x) multiplexing at up to three molecular targets. The use of an HRP-oligo integrates seamlessly into the SABER FISH workflow and only adds a brief (∼10 min) hydrogen peroxide incubation prior to sample permeabilization to inactive endogenous peroxidases and a short series of washes (∼10-30 mins) after development by HRP to remove residual HRP substrate. The use of the HRP-oligo and tyramide substrates is effectively cost-equivalent to performing conventional SABER, increasing the cost of visualization reagents per sample from ∼$0.10–$0.60 per sample (SABER) to $0.45–$2.30 per sample depending on the scale of the experiment (Table 1); in both cases, the cost of these development reagents is a minor fraction of the overall cost of the SABER/pSABER ISH experiment. We have demonstrated that pSABER: a) is readily compatible with colorimetric detection and histopathological stains, b) can substantially amplify target signals >20x beyond conventional SABER while maintaining expected subcellular localization patterns, c) can be multiplexed via solution exchange, d) is compatible with a broad range of optical systems including those commonly found in clinical labs, e) and can reveal detailed information about the spatial distribution of nucleic acid targets even on low numerical aperture and low magnification platforms. Given the simplicity, robustness and flexible readout options of the pSABER approach, we anticipate that it will serve as an enabling imaging technology in a wide set of clinical and research settings.

**Table 1.** Cost comparison of visualization reagents for SABER and pSABER. The cost per slide (center) and per imaging chamber slide well (right) are given for the key reagents needed to visualize SABER and pSABER.

| Reagent | Cost per slide (25 μl) | Cost per well (125 μl) |
| --- | --- | --- |
| Fluor-oligo (SABER) | \$0.11 | \$0.57 |
| HRP oligo (pSABER) | \$0.44 | \$2.22 |
| Tyramide fluor (pSABER) | \$0.01 | \$0.07 |

## Methods

### Cell culture

K29Mes human kidney mesangial cells^30^ were obtained from M. Saleem (University of Bristol) and cultured in RPMI-1640 medium supplemented with 10% (vol/vol) FBS, 1x L-glutamine, 1x non-essential amino acids, ITS+ supplement, and 50 U/ml penicillin, and 50 µg /ml streptomycin. K29Mes cells were grown at 33°C (permissive temperature) and were then shifted to 37°C (nonpermissive temperature) before fixation causing degradation of the temperature sensitive SV40 T-antigen to promote growth arrest and differentiation. Prior to experiments, cells were either seeded on 22 × 22 mm #1.5 coverslips in 6-well plates at a density of about 100,000 cells per well or seeded on Ibidi 8-well chamber slides at a density of about 70,000 and kept at 37°C overnight to allow adherence to coverslips. Cells were then rinsed in 1x PBS, fixed in either 4% (wt/vol) paraformaldehyde in 1x PBS or 10% (vol/vol) neutral buffered formalin for 10 minutes at room temperature, then rinsed twice in 1x PBS. EY.T4^34^ mouse embryonic fibroblast cells mouse embryonic fibroblast cells were grown in DMEM (ThermoFisher 10566016) supplemented with 15% (vol/vol) FBS (ThermoFisher A3160502), 50 U/ml penicillin, and 50 µg /ml streptomycin (ThermoFisher 15140122) 37°C in the presence of 5% CO_2_.

### Tissue

All deidentified human kidney tissue samples used in experiments were from the renal cortex region of the kidney obtained through nephrectomies performed at the University of Washington hospitals with local institutional review board approval (STUDY00001297). Fresh tissue was fixed for a minimum of 8 hours-overnight in 10% (vol/vol) neutral buffered formalin followed by rinsing in H_2_O and storage in 70% EtOH at 4°C till paraffin embedding. After embedding, 5 µm sections were obtained and placed on positively-charged microscope slides.

### FISH probes

Oligo probes targeting the human alpha satellite repeat, the human and mouse ITS1 RNA, the mouse 7SK RNA, and the mouse minor satellite repeat were purchased as individually column-synthesized DNA oligos from Integrated DNA Technologies. Probe sets targeting the human *ADAMTS1* mRNA, the human *UMOD* mRNA, and the mouse *Cbx5* mRNA were designed using PaintSHOP^31^ and ordered in plate oligo or oPool format from Integrated DNA Technologies. The complex oligo library used to target Xp22.32 was purchased from Twist Biosciences and amplified and processed as described previously^31^. The sequences of the oligos probes used are listed in Supplementary Dataset 1.

### Primer exchange reaction

To extend FISH probes into PER concatemers, 100 µl reactions were set up containing 1x PBS, 10mM MgSO4, 300 µM dNTP mix (dA, dC and dT only), 100 nM Clean.G hairpin, 40-600 U/mL of Bst LF polymerase (New England Biolabs), 500 nM of hairpin, and water to 90 µl.

Reactions were incubated for 15 minutes at 37°C, then 10 µl of 10 µM of primer was added to each reaction and incubated at 37°C for 2 hours, followed 20 minutes at 80°C to heat-inactivate the polymerase. To check the success of extensions and concatemer lengths, samples (diluted in 2x Novex TBE-Urea sample buffer) and DNA ladders (Low Range and 100 bp or 1 kb Plus, Thermo Fisher Scientific) were boiled at 95°C for 5 minutes then were run on 10-15% TBE-Urea gels for 30-60 minutes at 70°C using a circulating water bath hooked up to the gel cassette. Reactions were dehydrated via vacuum evaporation for long-term storage (at room temperature) and resuspended in hybridization solutions for downstream F/ISH experiments.

### Colorimetric DNA pSABER in fixed tissue culture cells

Fixed samples were permeabilized in 1x PBS with 0.1% (vol/vol) Triton X-100 for 10 minutes at room temperature, rinsed in 1x PBS, incubated in 0.5% (vol/vol) H_2_O_2_ in 1x PBS for 10 minutes, rinsed twice with 1x PBS, treated with 0.1N HCl for 5 minutes and washed twice with 2x SSC plus 0.1% (vol/vol) Tween-20 (2x SSCT). Coverslips were incubated in 2x SSCT with 50% (vol/vol) formamide for 2 minutes at room temperature and again for 20 minutes at 60°C. Samples were then inverted onto 25 µL of primary hybridization solution made up of 2x SSCT, 0.1% (vol/vol) Tween-20, 50% (vol/vol) formamide, 10% (wt/vol) dextran sulfate, 400 ng/µl of RNaseA, and 92 µl of dried PER reaction. Coverslips were placed in a humidified chamber, denatured at 78°C on a water-immersed heat block 3 minutes and then transferred to a benchtop air incubator for overnight incubation at 45°C. After hybridization with primary probes, samples were rinsed quickly in pre-warmed 2x SSCT before 4 rounds of 5-minute washes at 60°C and then 2 rounds of 2-minute washes in 2x SSCT at room temperature. For secondary hybridization (colorimetric detection), samples were rinsed in 1x PBS, incubated in blocking solution (1% wt/vol BSA and 0.1% vol/vol Tween-20 in 1x PBS), washed with 1x PBS twice, and inverted onto 25 µl of secondary hybridization solution consisting of 2x SSCT, 0.1% (vol/vol) Tween-20, 30% (vol/vol) formamide, 10% (wt/vol) dextran sulfate, and 100 nM HRP-oligo held at 37°C for 1 hour. Following secondary hybridization, coverslips were washed twice for 5 minutes in 1x PBS with 0.1% (vol/vol) Tween-20 at 37°C followed by a 5-minute wash in 1x PBS at room temperature. To develop color, samples were inverted onto 25 µl of DAB solution (1 drop of DAB Chromogen plus 50 drops of DAB substrate—Abcam ab64238) for 30 minutes, washed twice with tap water, counterstained with hematoxylin for 1 minute, rinsed with tap water twice, washed with 0.002% (vol/vol) ammonia water, and then washed in 70%, 95% and 100% (vol/vol) ethanol for a 3–5 mins each. To mount coverslips, they were each dipped in xylene 5 times and inverted onto a microscope slide holding a few drops of CytoSeal-XYL mounting medium. Mounted coverslips were kept in a microscope slide case at room temperature prior to imaging.

### Colorimetric DNA pSABER on metaphase spreads

Carrying out colorimetric DNA pSABER on human metaphase chromosome spreads was similar to the ‘Colorimetric DNA pSABER in fixed tissue culture cells’ protocol except for the steps taken right before primary probe hybridization. Slides with metaphase spreads were taken out of the −20°C freezer and denatured in 2x SSCT + 70% (vol/vol) formamide at 70°C for 3 minutes, followed by a 5-minute wash in ice-cold 70% (vol/vol) ethanol, in 90% (vol/vol) ethanol and in 100% ethanol and then allowed to air-dry before applying hybridization solution.

### Colorimetric RNA pSABER in fixed tissue culture cells

Fixed samples were permeabilized in 1x PBS with 0.1% (vol/vol) Triton X-100 for 10 minutes at room temperature, rinsed in 1x PBS, incubated in 0.5% (vol/vol) H_2_O_2_ in 1x PBS for 10 minutes, rinsed twice with 1x PBS, washed once with 1x PBS containing 0.1% (vol/vol) Tween (PBST), and once with 2x SSCT. Samples were then inverted onto 25 µl of primary hybridization solution made up of 2x SSCT, 0.1% (vol/vol) Tween-20, 10-40% (vol/vol) formamide, 10% dextran sulfate, and 92 µl of dried PER reaction. Coverslips were placed in a humidified chamber, denatured at 60°C on a water-immersed heat block for 3 minutes and then transferred to a benchtop air incubator for overnight incubation at 42°C. After hybridization with primary probes, samples were rinsed quickly in pre-warmed 2x SSCT before 4 rounds of 5-minute washes at 60°C and then 2 rounds of 2-minute 2x SSCT washes at room temperature. For secondary hybridization (colorimetric detection), samples were rinsed in 1x PBS, incubated in blocking solution (1% wt/vol BSA and 0.1% vol/vol Tween-20 in 1x PBS), washed with 1x PBS twice and inverted onto 25 µl of secondary hybridization solution consisting of 2x SSCT, 0.1% (vol/vol) Tween-20, 30% (vol/vol) formamide, 10% (wt/vol) dextran sulfate and 100nM HRP-oligo and held at 37°C for 1 hour in the dark. Following secondary hybridization, coverslips were washed twice for 5 minutes in PBST at 37°C followed by a 5-minute wash in 1x PBS at room temperature. To develop color, samples were inverted onto 25 µl of DAB solution (1 drop of DAB Chromogen plus 50 drops of DAB substrate for 30 minutes, washed twice with tap water, counterstained with hematoxylin for 1 minute, rinsed with tap water twice, washed with 0.002% (vol/vol) ammonia water, and then washed in 70%, 95% and 100% (vol/vol) ethanol for a few minutes each. To mount coverslips, they were each dipped in xylene 5 times and inverted onto a microscope slide holding a few drops of CytoSeal-XYL mounting medium for. Mounted coverslips were kept in a microscope slide case at room temperature prior to imaging.

### Colorimetric RNA pSABER in FFPE tissue

To deparaffinize FFPE sections, slides were placed in a slide rack and submerged in xylene twice for five minutes, incubated in 100% ethanol twice for 1 minute then removed and allowed to be air dried for 5 minutes at room temperature. To prepare FFPE tissue sections for *in situ* hybridization, pretreatment reagents from Advanced Cell Diagnostics (ACD) were used. Sections were incubated in 5–8 drops of ACD’s hydrogen peroxide for 10 minutes at room temperature and rinsed with distilled water twice. The tissue slides were then submerged into the boiling 1x Target Retrieval solution from ACD for 10 minutes and immediately transferred to a staining dish containing distilled water, followed by another water rinse. Slides were washed in 100% ethanol quickly and allowed to be air dried. Lastly, sections were incubated with ACD’s Protease Plus, placed in a humidified chamber inside an air incubator at 40°C for 40 minutes and washed with distilled water twice. Pretreatment steps were followed by primary probe hybridization and all the following steps detailed in the protocol for ‘Colorimetric RNA pSABER in fixed tissue culture cells’ protocol.

### Fluorescent RNA pSABER in fixed tissue culture cells

Fixed cells that were grown on Ibidi 8-well chamber slides underwent the same steps described in the ‘Colorimetric RNA pSABER in fixed tissue culture cells’ protocol up to the wash prior to undergoing secondary hybridization. For the secondary hybridization 125 µl of secondary hybridization solution consisting of 2x SSCT, 0.1% (vol/vol) Tween-20, 10% (wt/vol) dextran sulfate and 100nM HRP-oligo was added to the samples and held at 37°C for 1 hour in the dark. Cells in each well were then washed 3 times for 5 minutes in PBST at 37°C. To employ signal amplification via tyramide, fluorescent tyramide reagents that were purchased from Tocris were optimized and final concentrations of 480nM, 1.1 µM, and 3.75 µM for fluorescein, cy3 and cy5 tyramides were established respectively. To develop signal, 125 µl of tyramide solution consisting of 1x PBS, 0.0015% (vol/vol) H_2_O_2_, and one of the fluorescent tyramides was added to the samples for 10 minutes at room temperature. To quench enzyme activity, samples were washed 3 times for 5 minutes with a quenching solution containing 10 mM sodium azide and 10 mM sodium ascorbate in PBST and 2 extra washes with PBST alone. Prior to being imaged, 125 µl of SlowFade Gold Antifade Mountant with DAPI was added to each well.

### DNA SABER-FISH and Fluorescent DNA pSABER in fixed tissue culture cells

DNA FISH was performed essentially as described previously^28^. Cells were grown on Ibidi 18-well chamber slides at a seeding density of ∼7000 cells/ well in a mammalian tissue culture incubator and fixed in 4% (wt/vol) paraformaldehyde for 10 minutes. The samples were permeabilized in 1x PBS + 0.5% (vol/vol) Triton x-100 for 10 minutes at room temperature (hereafter RT), rinsed in 1x PBS + 0.1% Tween-20 (hereafter PBS-T) for 5 minutes, incubated in 0.1N HCl for 5 minutes, rinsed with PBS-T, incubated in 3% (vol/vol) H_2_O_2_ in PBS-T for 10 minutes (pSABER only), rinsed twice with 2x SSC + 0.1% (vol/vol) Tween-20 (hereafter 2x SSCT), incubated in 2x SSCT + 50% (vol/vol) formamide for 2 minutes, incubated in 2x SSCT+ 50% (vol/vol) formamide at 60°C for 20 minutes on a water-immersed heat block, and denatured at 80°C for 3 minutes in a hybridization solution consisting of 2x SSC, 0.1% (vol/vol) Tween-20, 50% (vol/vol) formamide, 10% (wt/vol) dextran sulfate, 0.4ug/uL RNase A, and 100 pmol of unpurified PER reaction. The slide was then held at 37°C overnight in a humidified chamber. On the next day, 4 washes’ worth of 2x SSCT were pre-warmed to 60°C and the cells were rinsed twice with RT 2x SSCT. The cells were then rinsed 4x with the pre-warmed 2xSSCT on a 60° heat block, with each rinse taking 5 minutes. This was followed by two quick RT rinses with 2x SSCT and three quick rinses in 1x PBS. The cells were then incubated at 37°C for 1 hour in a secondary hybridization solution consisting of 0.1uM fluor oligos (SABER) or HRP-oligos (pSABER) in 1x PBS. During this hour, three washes’ worth of PBS-T were pre-warmed to 37°C. After the secondary hybridization was complete, the cells were rinsed 3x in the pre-warmed PBS-T, with each wash taking 5 minutes. For pSABER, cells were then incubated for 10 minutes at RT in a labeling solution consisting of fluorescein tyramide (Torcis 6456) diluted 1:50,000 from its 24 mM stock and 0.0015% (vol/vol) H_2_O_2_ in PBS-T. pSABER development, the cells were rinsed 3x for 5 minutes each in a reaction quenching solution consisting of 10mM sodium ascorbate and 10mM sodium azide in PBS-T. Finally, the cells were rinsed twice in PBS-T and SlowFade Gold antifade mountant with DAPI was added.

### Fluorescent RNA pSABER in FFPE tissue

FFPE tissue sections underwent the same deparaffinization and pretreatment steps detailed in the protocol for Colorimetric RNA pSABER in FFPE tissue. Following pretreatment incubations, tissue samples went through the same stages laid out in the protocol for ‘Fluoresecent RNA pSABER in fixed tissue culture cells’, however, they did not get SlowFade Gold Antifade Mountant with DAPI at the end. Prior to imaging, tissue slices were stained with 0.1 ug/mL DAPI (in 1x PBS) for 5 minutes and washed with 1x PBS. To reduce autofluorescence, the samples were then treated with 1x TruBlack Lipofuscin Autofluorescence Quencher (in 70% ethanol) for 1 minute taking care not to dry the tissue sections. Slides were washed 3 times with 1x PBS and coverslips were mounted using Prolong Gold Antifade Mountant.

### Exchange RNA pSABER in fixed tissue culture cells

The steps are identical to the steps of the ‘Fluorescent RNA pSABER in fixed tissue culture cells’ protocol, but instead of adding the mountant with DAPI at the end, the samples were washed with 60% (vol/vol) formamide in PBST for 15 minutes at room temperature to remove the first round of HRP-oligo, followed by 2 washes with 1x PBS for 1 minute, another 2 washes for 2 minutes and a final 5-minute wash. To enable another round of detection, samples were incubated with BSA blocking solution again, and got 125 µl of the second HRP-oligo hybridization solution followed by all the steps detailed in the ‘Fluorescent RNA pSABER in fixed tissue culture cells’ protocol that came after secondary hybridization. At the end either another round of detection was performed or 125 µl of SlowFade Gold Antifade Mountant with DAPI was added to each well prior to imaging. The following three tyramides were used: fluorescein at 1:50,000 from the 24 mM stock (Tocris 6456); Cy3 at 1:15,0000 from the 16 mM stock (Tocris 6457); Cy5 (Tocris 6458) at 1:4,000 from the 15 mM stock.

### Microscopy

Transmitted light microscopy of colorimetric pSABER samples was performed on an Olympus BX41 upright microscope using 20x Plan N.A. 0.25, 40x UPlan N.A. 0.75 or 60x Ach N.A. 0.80 air objectives with images captured using a Leica DFC420 camera. Confocal imaging of fluorescent SABER and pSABER samples was performed using Yokogawa CSU-W1 SoRa spinning disc confocal attached to a Nikon Eclipse Ti-2 microscopy. Excitation light was emitted at 30% of maximal intensity from 405 nm, 488 nm, 561 nm, or 640 nm lasers housed inside of a Nikon LUNF 405/488/561/640NM 1F commercial launch. Laser excitation was delivered via a single-mode optical fiber into the into the CSU-W1 SoRa unit. Excitation light was then directed through a microlens array disc and a ‘SoRa’ disc containing 50 µm pinholes and directed to the rear aperture of a 100x N.A. 1.49 Apo TIRF oil immersion objective lens by a prism in the base of the Ti2. Emission light was collected by the same objective and passed via a prism in the base of the Ti2 back into the SoRa unit, where it was relayed by a 1x lens (conventional imaging) or 2.8x lens (super-resolution imaging) through the pinhole disc and then directed into the emission path by a quad-band dichroic mirror (Semrock Di01-T405/488/568/647-13×15×0.5). Emission light was then spectrally filtered by one of four single-bandpass filters (DAPI: Chroma ET455/50M; ATTO 488: Chroma ET525/36M; ATTO 565: Chroma ET605/50M; Alexa Fluor 647: Chroma ET705/72M) and focused by a 1x relay lens onto an Andor Sona 4.2B-11 camera with a physical pixel size of 11 µm, resulting in an effective pixel size of 110 nm (conventional) or 39.3 nm (super-resolution). The Sona was operated in 16-bit mode with rolling shutter readout and exposure times of 70–300 ms. Automated slide-scanning microscopy was performed on a Keyence BZ-X800. Excitation light from the integrated white light source was emitted at 25% of maximal intensity, passed through the ‘ET DAPI’, ‘ET GFP’, ‘Cy3’, or ‘Cy5’ filter cube for excitation band filtering and was directed to the sample by a 10x N.A. 0.45 Plan Apo, 20x N.A. Plan Apo, or 40x N.A. 0.95 air objective lens. Emission light was collected by one of the objective lenses and directed into the microscope’s emission path by one of the filter cubes, after which it was spectrally filtered while exiting the filter cube and focused by a 1x relay lens onto an integrated CCD camera with a physical pixel size of 7.549 µm, resulting in effective pixels sizes of 188.7 nm (40x), 377.4 nm (20x), or 754.9 nm (10x). The CCD camera was operated in 14-bit mode without pixel binning and exposure times of 16.7–666.7 ms. Images were processed in ImageJ/Fiji^36,37^ and Adobe Photoshop.

### Quantitative analysis of peak puncta intensities

We developed an image analysis pipeline to detect and segment puncta from both SABER and pSABER experiments quantify their peak intensities. The pipeline is capable of loading .nd2 image files containing channels with a nuclear stain (here DAPI) and fluorescent puncta, segmenting nuclei in 2D and detecting puncta inside the nuclei, and returning raw intensity values for each punctum. Each image is first loaded with the nd2reader python package, and DAPI / GFP channels are split for separate processing pathways. For the DAPI channel, 2D segmentation was performed with the python package StarDist, specifically the “2D_versatile_fluo” pretrained model^38,39^. To generate more accurate segmentation results, nuclear segmentation was performed on a max-intensity projection which was downsampled by a factor of 16. The resulting segmentation mask was then upsampled to the original resolution. For the GFP channel, a maximum-intensity projection was generated and spot detection was performed using the *blob_log* function from scikit-image^40^. Spot detection with blob_log consisted of fitting a 2D Gaussian distribution to each punctum and returned the highest-intensity pixel as the 16-bit intensity of the spot. For SABER, sigma parameters were set as follows: threshold = 0.02, min_sigma=0.001, max_sigma=0.5, num_sigma=20. For pSABER, sigma parameters were set as: min_sigma=0.01, max_sigma=1, num_sigma=20. Detected spots were then filtered based on overlap with the nuclear segmentation mask. Data from each condition was aggregated and visualized with matplotlib. To quantify signal amplification, the cumulative distribution function for the SABER and pSABER spot intensities was generated, and the distributions were normalized to the SABER condition (median = 1). Signal amplification here was defined as the resulting median of the pSABER distribution.

## Code and Data Availability

The source code for the automated analysis of peak puncta intensities as well as example input images is available at https://github.com/beliveau-lab/SABER-Spot-Quant under a MIT license. Additional primary microscopy data will be made available upon request.

## Author Contributions

S. Attar, S. Akilesh, and B.J.B. conceived the study. V.A.B. wrote and optimized software code. S. Attar, V.A.B., E.K.N., and A.F.T. performed experiments. S. Attar, V.A.B., S. Akilesh, and B.J.B. wrote the manuscript. All authors edited and approved the manuscript. D.M.S., J.S., J.A.L., S. Akilesh, and B.J.B. supervised the work.

## Acknowledgements

The authors thank members of the Shechner, Shendure, Lieberman, Akilesh, and Beliveau labs for helpful discussion about this work. We thank N. Peters of the UW W.M. Keck Microscopy Center and D. Fong of Nikon for assistance with microscopy during the development of pSABER and S. Jiang for helpful advice about working with HRP in tissues. This work was supported by a Damon Runyon Dale F. Frey Breakthrough Award (32-19 to B.J.B.), the National Institutes of Health (under grants 1R35GM137916 to B.J.B, 1UM1HG011586 to J.S. and B.J.B., 1R01DK130386 to S. Akilesh, 1R01GM138799-01 to D.M.S.), the Andy Hill Cancer Research Endowment (under a COVID-19 Response Grant Award to B.J.B. and S. Akilesh), and the Diabetic Complications Consortium (under grant 19AU3987 to S. Akilesh and B.J.B.). This project was also supported in part by a Building Bridges Award to J.A.L. and S. Akilesh from the Department of Laboratory Medicine and Pathology. A.F.T. was supported by NIH training grant T32GM007750.

---

**Fig. S1.**
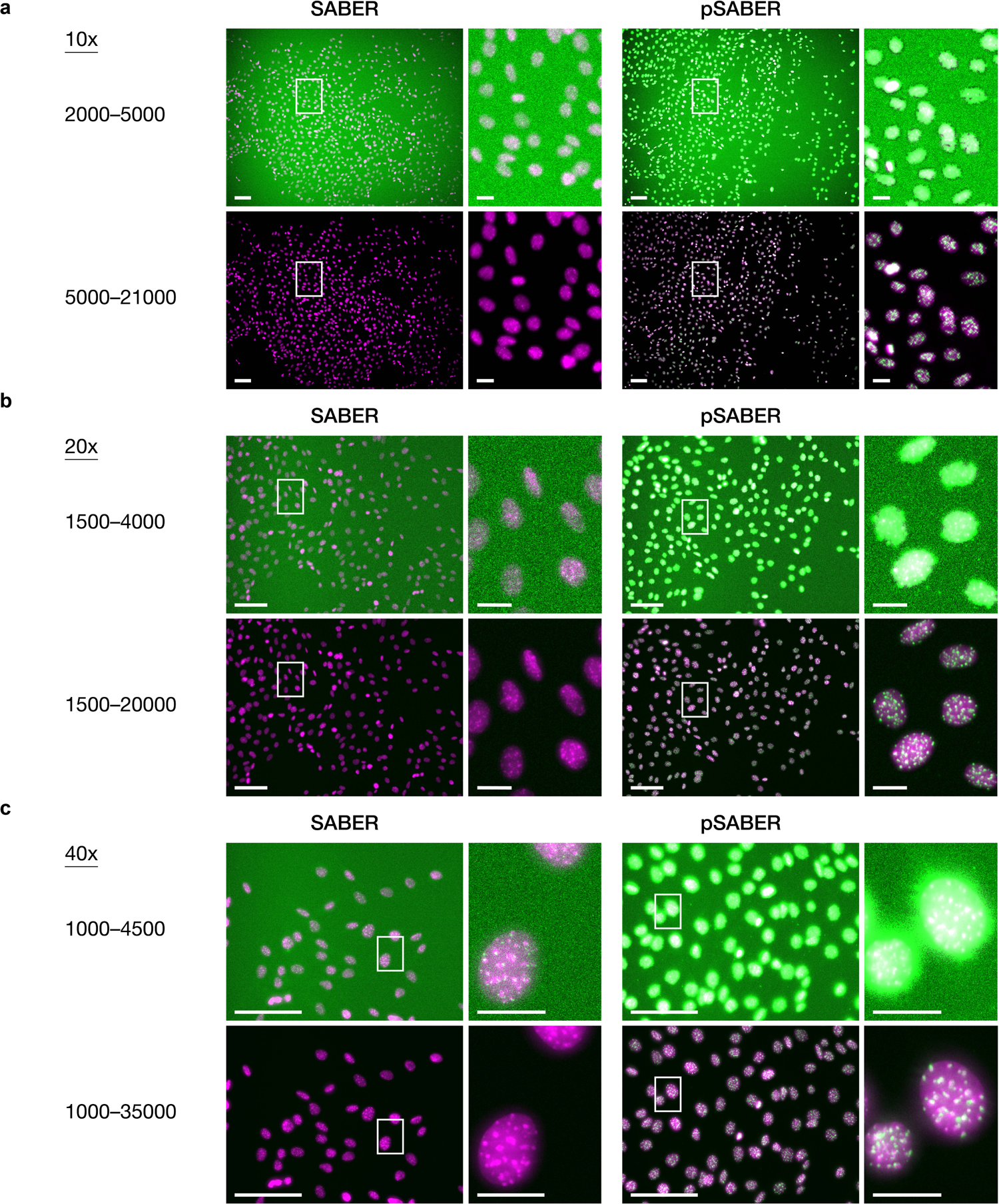
Comparison of SABER and pSABER on a slide-scanning microscope. **a**, SABER (left) and pSABER (right) visualization of mouse major satellite DNA imaged using a 10x air objective. **b**, SABER (left) and pSABER (right) visualization of mouse major satellite DNA imaged using a 20x air objective. **c**, SABER (left) and pSABER (right) visualization of mouse major satellite DNA imaged using a 10x air objective. Each image is presented at the indicated high (top) and low (bottom) contrast settings to allow for visual comparisons between signals occupying different portions of the 16-bit dynamic range (0–65535). Images are maximum projections in Z. Scale bars, 20 µm (insets) or 100 µm (fields of view).

